# Finding a suitable library size to call variants in RNA-seq

**DOI:** 10.1101/2019.12.18.881870

**Authors:** Anna Quaglieri, Christoffer Flensburg, Terence P Speed, Ian J Majewski

## Abstract

**Background:** RNA-Seq allows the study of both gene expression changes and transcribed mutations, providing a highly effective way to gain insight into cancer biology. When planning the sequencing of a large cohort of samples, library size is a fundamental factor affecting both the overall cost and the quality of the results. While several studies analyse the effect that library size has on differential expression analyses, sensitivity analysis for variant detection has received far less attention.

**Results:** We simulated shallower sequencing depths by downsampling 45 AML samples that are part of the Leucegene project, which were originally sequenced at high depth. We compared the sensitivity of six methods of recovering validated mutations on the same samples. The methods compared are a combination of three popular callers (MuTect, VarScan, and VarDict) and two filtering strategies. We observed an incremental loss in sensitivity when simulating libraries of 80M, 50M, 40M, 30M and 20M fragments, with the largest loss detected with less than 30M fragments (below 90%). The sensitivity in recovering indels varied markedly between callers, with VarDict showing the highest sensitivity (60%). Single nucleotide variant sensitivity is relatively consistent across methods, apart from MuTect, whose default filters need adjustment when using RNA-Seq. We also analysed 136 RNA-Seq samples from the TCGA-LAML cohort, assessing the change in sensitivity between the initial libraries (average 59M fragments) and after downsampling to 40M fragments. When considering single nucleotide variants in recurrently mutated myeloid genes we found a comparable performance, with a 3% average loss in sensitivity using 40M fragments.

**Conclusions:** Between 30M and 40M fragments are needed to recover 90%-95% of the initial variants on recurrently mutated myeloid genes. To extend this result to another cancer type, an exploration of the characteristics of its mutations and gene expression patterns is suggested.

## Background

RNA-Seq is routinely used to quantify transcripts, detect fusion genes and differential splicing. It can also be used to call mutations, a key component in the study of cancer genomes. This makes RNA-Seq a cost effective choice in cancer research. However, calling variants in RNA-Seq cancer samples is often overlooked due to the large number of possible sources of bias. We are only able to call variants if they belong to transcribed DNA, and gene expression variation causes the transcriptome-wide depth to be highly heterogeneous. The variant allele frequency (VAF) in cancer samples can also be affected by other sources of variation, such as normal tissue contamination, differing clonality and copy number changes. When working with human material we are often limited by the number of samples available, and this makes decisions about the sequencing depth (or library size) especially critical. Previous research has highlighted the importance of sequencing depth in many fields of genome research, including transcriptome sequencing, but typically this has considered differential expression (DE) analysis, transcript discovery and differential splicing[1]. In that study a staged sequencing approach was presented as a useful tool for determining the parameters of the sequencing experiment (e.g. the number of replicates or the number of mapped reads). Numerous papers have been published around the power to detect DE genes in RNA-Seq, but the discussion has mainly concerned the number of replicates needed and the statistical tools applied[2, 3, 4]. The number of transcripts that can be identified and the number of potentially false positive DE genes increases steadily as sequencing depth increases[5]. Earlier work on the power to detect DE genes based on the number of replicates, sequencing depth and analytical tools used suggests that the number of replicates is more important than the read depth, and that going beyond 20M fragments does not increase the power[6]. Sensitivity analysis for variant calling from RNA-Seq data has received less attention. A staged sequencing approach has previously been adopted to study the sensitivity of SNP calling with whole exome sequencing (WES) and whole genome sequencing (WGS) in normal tissues, using HapMap SNPs as gold standard[7]. A mean-on-target of 40X reads was found to be enough to reach 95% sensitivity, but it was not clear how to decide on the total library size. The most recent study on the sensitivity of SNP calling in RNA-Seq in cancer samples, offers an extensive overview of the use of different combinations of aligners and callers[8]. That study found that with all methods, the sensitivity remains >90% with a total read depth of >10X at the variant site, and concluded that the proportion of variants recovered by RNA-Seq will more likely depend on the total depth at the variant site than on the total library size. However, the total depth distribution depends on the expression level of the corresponding gene, and this is proportional to the total library size. The motivation for the present study was the need to inform the in-house sequencing of a cohort of Core Binding Factor Acute Myeloid Leukaemia (CBF-AML) RNA-Seq samples in order to allow accurate DE analysis and variant calling. Given the lack of available literature regarding the library size needed to reach an adequate sensitivity in cancer RNA-Seq variant calling, we developed a staged sequencing approach using 45 publicly available and deeply sequenced CBF-AML RNA-Seq samples[9], where mutations that were detected in the RNA were validated on matched DNA using targeted Sanger Sequencing if they were not present in the COSMIC database[10]. We used this set of validated variants in order to get estimates of the sensitivity at shallower depths and validated our findings on a larger independent AML cohort.

## Results

### Sensitivity in the Leucegene cohort

We used 45 CBF-AML RNA-Seq samples that were deeply sequenced with 100 base pair (bp) paired end (PE) reads to compute the sensitivity in recovering 88 validated mutations at lower levels of sequencing depth[9] (Table 1, Figure S1). This was done by simulating smaller library sizes by random downsampling of the reads in the initial samples. We will refer to these validated mutations as the *truth set*. After alignment, the initial samples have a mean of 113M mapped PE reads (min 77.8M - max 187.4M, 93.4% mean mapping rate).

**Table 1:** Type of variants in the Leucegene truth sets. Variants used as the truth set were previously validated in a set of 45 CBF-AML RNA-Seq samples[9]. Variant types are inferred from the information in the published study and by the variant calls performed on the initial samples. A short indel (insertion/deletion) is an indel <10bp long; composite indels are mutations including both inserted and deleted nucleotides; SNVs are single nucleotide variants.

| Mutation type | Min VAF | Mean VAF | Max VAF | N |
| --- | --- | --- | --- | --- |
| Composite indel | 0.06 | 0.25 | 0.56 | 9 |
| Long insertion | 0.41 | 0.5 | 0.64 | 3 |
| Short deletion | 0.09 | 0.24 | 0.38 | 2 |
| Short insertion | 0.07 | 0.33 | 0.84 | 15 |
| SNVs | 0.05 | 0.37 | 0.97 | 58 |
| Indel-Not reported | 0.84 | 0.84 | 0.84 | 1 |

We downsampled the initial unaligned files at five fixed library sizes of 80, 50, 40, 30, and 20 million PE reads (or fragments) to simulate shallower depths. Two samples have library size marginally below 80M and we used all the reads as the initial run. At each library size, the random downsampling was replicated more than once to account for downsampling variability (see ‘Downsampling strategy’ section in Methods). At every stage we called variants using three different callers, previously used to call variants in RNA-Seq: MuTect2[11], VarScan2[12], and VarDict[13]. We will refer to MuTect2 as MuTect and to VarScan2 as VarScan. Figure 1 shows the sensitivity in recovering all the variants in the truth set. The sensitivity is computed using two different filtering strategies: 1) *default-filters* which are based on each caller’s default settings and a set of variants detected in a panel of normals (PON) comprising RNA-Seq samples from CD34+ cells; 2) *annotation-filters* which uses external databases, the PON variants and other quality filters (see details in ‘Variant filtering’ section in Methods).

**Figure 1:**
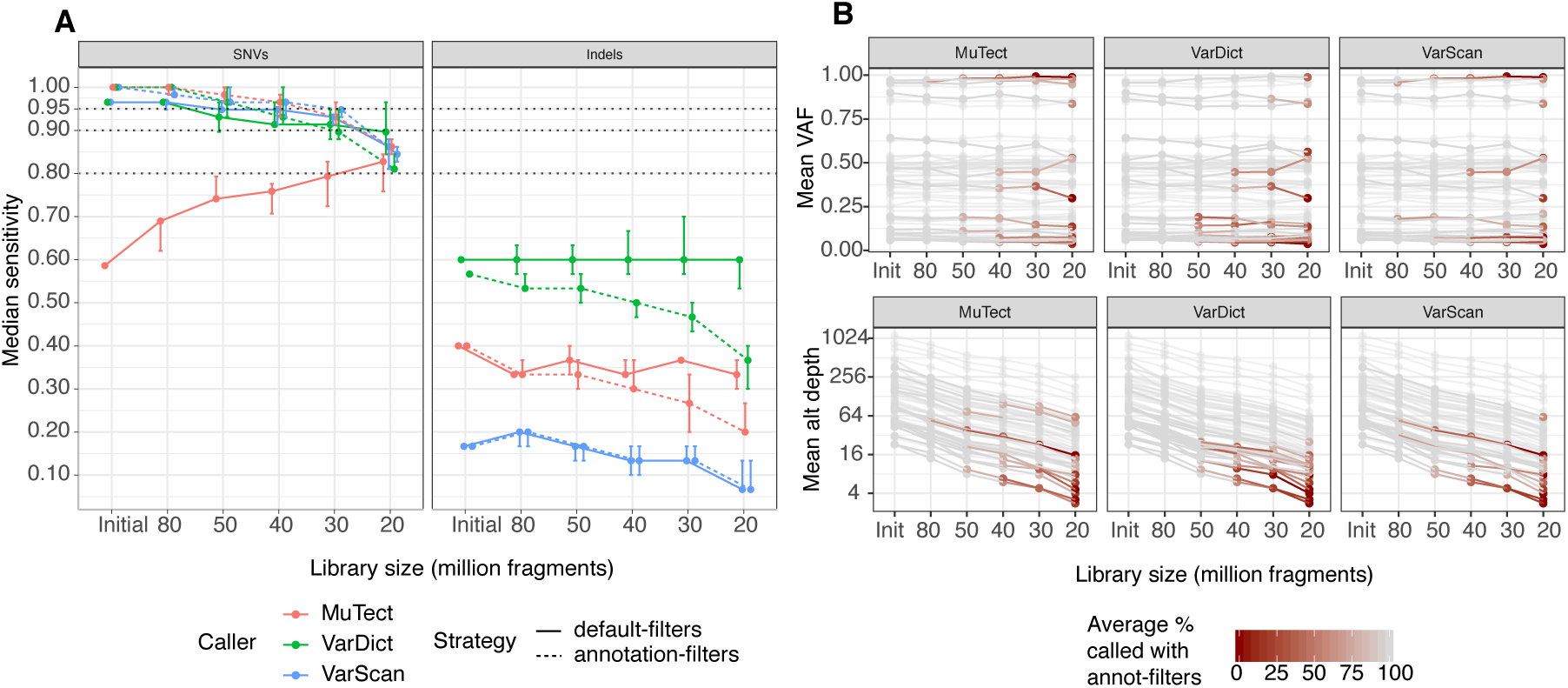
Sensitivity in recovering the variants in the truth set using the Leucegene RNA-Seq samples. **A**. Median with maximum and minimum sensitivity (vertical bars) for recovering the SNVs (left plot) and indels (right plot) in Table 1, across random downsampling runs using different library sizes. Each estimated median sensitivity represents the median across 5 random downsamplings (only 3 for 80M libraries) of the initial RNA-Seq libraries at a specific library size. The solid lines are the sensitivities obtained using the default-filters and the dotted lines are obtained with the annotation-filters. **B**. Average VAF (top plot) and alternative depth (bottom plot) on the log scale at a variant site for the variants in the truth set using different library sizes. A line in each plot represents one mutation in the truth set. Each dot is coloured according to the average number of times a variant was called by one caller using the annotation-filters across replicated downsampling runs at one specific library size.

The sensitivity for detecting single nucleotide variants (SNVs, Figure 1A left plot) is comparable across filtering strategies and callers apart from the unusual behaviour of MuTect with default-filters, where sensitivity increased as the library size decreased. MuTect behaviour is due to the `clustered events` filter which removes variants found on haplotypes where other variants are already detected (Figure S2). This behaviour was also observed in a previous study comparing variants called from matched RNA-Seq and WES samples where the same flag was responsible for filtering the largest number of RNA variants[14].

Using knowledge from external databases allows retention of two NRAS SNVs present in COSMIC[10], which are discarded by default-filters as they are also present in the PON samples at very low frequency. The SNV sensitivity remains above 95% for all callers with the initial and 80M libraries and it incrementally decreases between the 50M and the 30M libraries, remaining around 90%. 100% of the initial variants are recovered with the initial and the 80M libraries using the annotation-filters. The largest drop in sensitivity is observed when moving from 30M to 20M fragments, where in some cases only 80% of the initial SNVs were recovered. With the default-filters, the median sensitivity for SNVs decreases by a maximum of 5% between the initial and the 30M libraries when using VarScan or VarDict, reaching a 10% loss with 20M fragments (Figure S3). The drop in sensitivity using subsequently smaller library sizes is larger when using the annotation-filters, even though the sensitivity with larger libraries is higher using this strategy. The sensitivity in recovering indels (Figure 1A right plot) varied markedly between callers, with VarDict calling consistently more indels than the other callers, but still only achieving a maximum of 60% recall. The large difference in indel sensitivity between callers is partly due to a bias in reporting indels. VarDict uses the same approach as the km[15] caller, used to create the truth set, and it is the only caller adopted here which was developed for both DNA and RNA. Indel sensitivity slightly increases with MuTect and VarScan if only a partial match with the allele in the truth set is required (Figure S4). For these reasons, the choice of a suitable library size will be based on SNV sensitivity, and indel sensitivity should be assessed using more appropriate and comparable callers. The unusual behavior of MuTect with default-filters is not observed for indels. This could be because a large number of indels are not detected by MuTect even with the initial samples. Therefore, the sensitivity curves do not appropriately reflect the change in the behaviour of the caller at different depths. The majority of the SNVs missed at shallower sequencing depths are either not reported by a caller or subsequently filtered by quality thresholds, especially when using annotation-filters (see flags of variant missed in Figure S5). The quality filters are mainly affected by the hard threshold on the total and alternative depth at a variant site (see details in ‘Annotation filters’ section in Methods).

The VAF of the variants remains stable across library sizes (Figure 1B top plot), while the alternative and total depths at a variant site decrease steadily (Figures 1B bottom plot, Figure S6, distribution of the initial alternative and total depth in Figure S7), with low alternative depth characterizing a large part of the variants lost. The median alternative depth of the variants missed by a caller has a sharp drop when considering less than 50M fragments. The alternative depth varies between 30X to 50X when using libraries larger than 50M fragments and between 5-9X for smaller libraries (Table S2). This is because the variants missed with larger libraries are not called due to presence in the PON, or mismatches with the alternative allele detected in the truth set. On the other hand, as the library size decreases, a larger number of variants present in smaller clones or lowly expressed loci, are lost. This is also observed in the bottom plot in Figure 1B, where more red lines (variants lost) are noticeable with less than 50M fragments.

### Sensitivity in the TCGA-LAML cohort using validated WGS and WES variants

Using the Leucegene samples we showed that there is limited sensitivity below 30M fragments. Increasing the library size >30M induces an incremental gain in sensitivity and with 40M PE all callers recover >90% of the initial variants. Therefore, we decided to analyse the loss in sensitivity in an independent AML cohort by using 136 50bp PE RNA-Seq samples from the TCGA-LAML cohort[16], choosing 40M fragments as the target library size. We analysed the loss in sensitivity between the initial libraries and the ones downsampled to 40M PE reads, where the initial samples have an average of 58.7M mapped reads (min 36.7M - max 69.6M) and a lower mapping quality than the Leucegene CBF-AML samples (75.6% mean mapping rate). The available BAM files were downsampled at different proportions depending on their mapping rate, in order to obtain a number of mapped reads similar to that obtained with the 40M downsampled Leucegene libraries (see ‘Downsampling strategy’ section in Methods). We called variants with VarDict, MuTect and VarScan on the initial and downsampled BAM files, and evaluated the sensitivity in recovering SNVs from two truth sets created from the list of validated WES and WGS variants[16]. The truth sets are: *Set1*, including 1,643 SNVs from the published variants after removing intergenic and intronic SNVs (Figure 2A left plot); *Set2*, which is a subset of Set1 including 169 SNVs from recurrently mutated myeloid genes (Figure 2A right plot, Table S3, details in ‘TCGA truth sets’ section in Methods). Both the default-filters and the annotation-filters were used for variant filtering with some minor exceptions from the Leucegene analysis (see ‘TCGA truth sets’ section in Methods). Due to the challenges induced by different indel representation across callers described in the previous section, we only assessed SNVs sensitivity.

**Figure 2:**
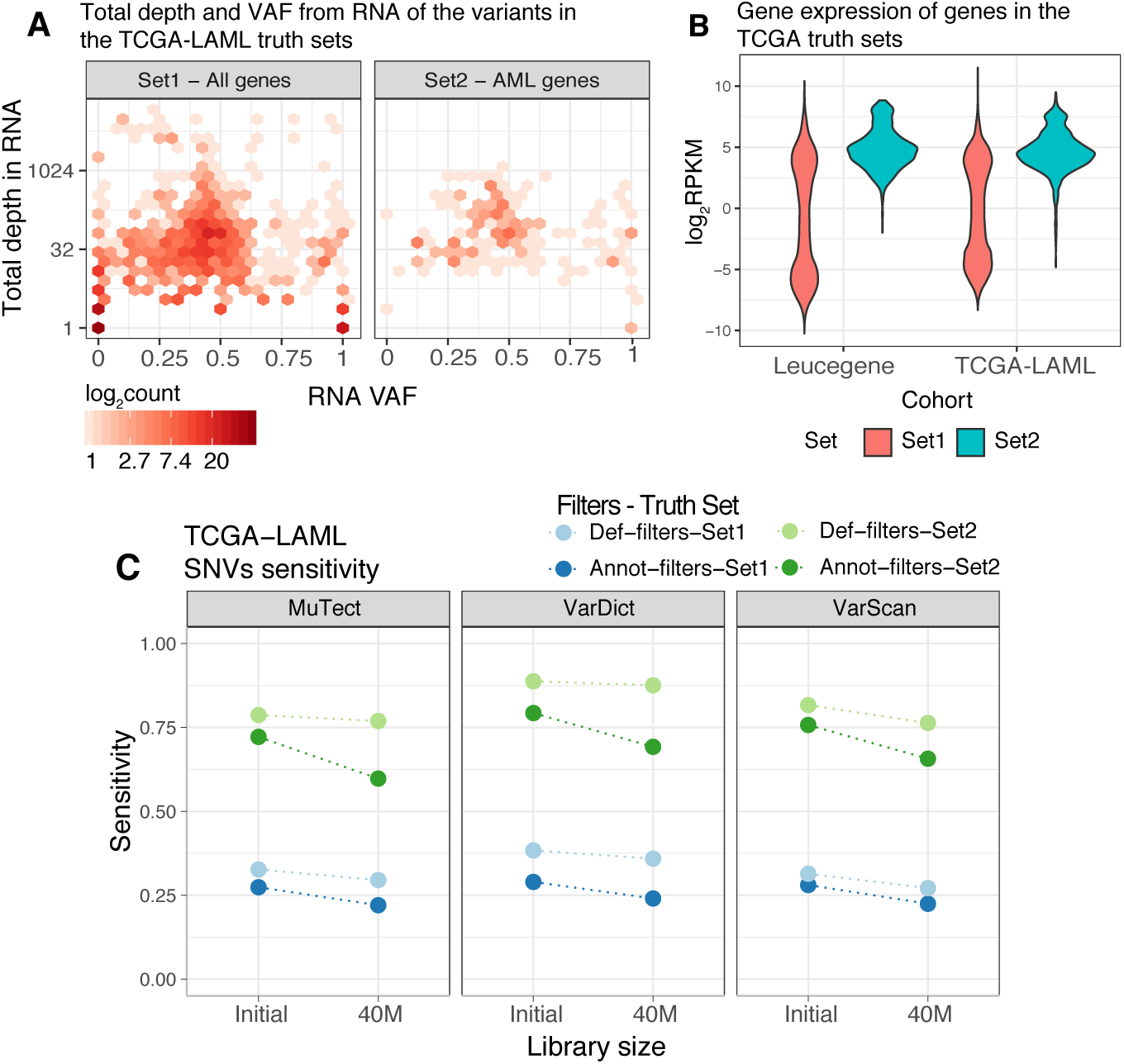
Sensitivity using the TCGA-LAML truth sets. **A**. 2D Density plots of the RNA VAF (x-axis) against the total depth on a logarithmic scale (y-axis) of the SNVs in the validated sets of variants: Set1 on the left and Set2 on the right. **B**. Violin plots of the log_2_RPKM of the genes with variants detected in Set1 and Set2. The gene expression distributions are provided for the TCGA-LAML and the Leucegene libraries using the initial samples, before downsampling. **C**. Sensitivity in recovering SNVs from the TCGA-LAML cohort in the initial and downsampled libraries using a combination of the two truth sets, callers and filtering strategies.

Genes whose variants belong to Set2 tend to be more expressed than genes in Set1 in both the Leucegene and the TCGA-LAML cohorts (Figure 2B). This is also reflected at the variant level, where a large number of variants from Set1 has very low total depth at the variant site in the RNA samples (Figure 2A left plot), while when only restricting to myeloid genes, there is only a low number of variants with very low total depth (Figure 2A right plot). This implies that the genes commonly mutated in AML also tend to be more expressed. Figure 2C summarizes the sensitivity with the initial and the downsampled TCGA-LAML data. The default-filters always outperform the annotation-filters. As expected, the sensitivity in recalling the variants in Set1 (blue shaded dots) is quite low, ranging between 22% and 38% across callers, with marginal differences between library sizes, (average 4% decrease in sensitivity). The recall rate improves if only variants on recurrently mutated myeloid genes are considered (green shaded dots). VarDict with default-filters has the highest sensitivities, recovering 89% and 88% of the SNVs with the initial and the downsampled libraries respectively. Within this truth set, the filtering strategy used has a larger impact on the sensitivity. The sensitivity using default-filters remains largely stable across callers and library sizes, with an average decrease of 3% between the initial and the 40M libraries. However, a larger decrease in sensitivity is observed when applying the annotation-filters, reaching a 10% decrease. Many of the variants are filtered by the hard thresholds on the total and alternative depth at a variant site as well as because of their proximity to exon boundaries (see flags and alternative depth of missed variants in Figures S8).

Figure 3A offers a breakdown of the per gene sensitivities using default-filters and considering only genes whose variants are in Set2. MuTect’s poor performance on the top recurrently mutated genes, FLT3, IDH1, and IDH2 is again caused by the clustered events filter, discussed in the ‘Sensitivity in the Leucegene cohort’ section and sensitivity improves when applying annotation-filters (Figure S9). Both VarDict and VarScan recover almost all events on these genes which are highly expressed across the TCGA-LAML samples (log_2_RPKM > 3.5 across the three genes, log_2_RPKM > 4.5 for FLT3 and IDH2). VarDict is the caller with the highest sensitivity, reaching > 90% recovery rate with the top six mutated genes (DNMT3A, IDH2, RUNX1, IDH1, FLT3 and TP53) using both the initial and the 40M PE reads libraries. The genes with the lowest sensitivities are U2AF1, KDM6A, RAD21 and STAG2. Apart from U2AF1, not many variants are present on the other genes and several SNVs are filtered out due to low quality of the alternative alleles and low total depth. UA2AF1 has the lowest sensitivities across all genes in Set2, apart when using VarScan. Not surprisingly, this gene also has the lowest expression levels among all myeloid genes considered (log_2_RPKM < 2.5, Figure 3B). As the gene expression decreases there is an increased discordance between callers (red and darker shaded dots in Figure 3B) and only around 50% of the variants are detected on average with log_2_RPKM < 3.

**Figure 3:**
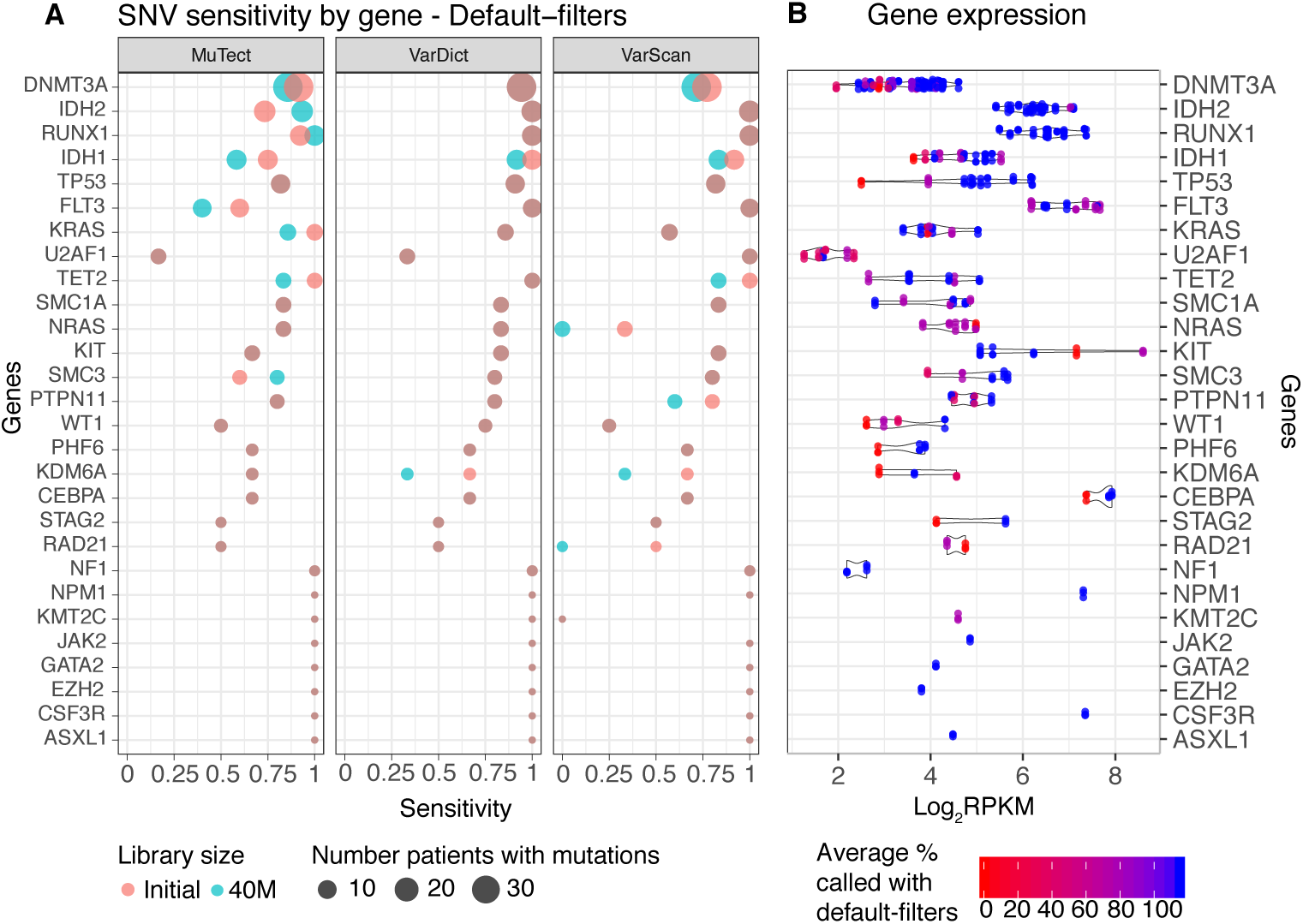
SNVs sensitivity and expression of the recurrently mutated AML genes using the TCGA-LAML cohort. **A**. Sensitivity in recovering mutations on recurrently mutated AML genes (Set2) using the TCGA-LAML cohort with callers default-filters. The size of the dots is proportional to the number of times a gene is mutated and genes were ordered by mutation load, with the most mutated genes at the top. Red dots corresponds to the results obtained with the initial library sizes and cyan dots using the downsampled libraries. **B**. Expression distribution of recurrently mutated myeloid genes across the TCGA-LAML RNA-Seq samples used. The genes are reported in the same order as in panel **A**. Each dot corresponds to a sample and dots are coloured based on the percentage of times the variants detected on a sample are called by the callers using default-filters and the 40M libraries. A patient can harbour more than one mutation per gene. Horizontal violin plots are drawn below the dots.

### Sensitivity by total depth at a variant site

We estimated the sensitivity as a function of the total depth at a variant site for both the TCGA-LAML samples using variants from Set1 (Figure 4A) and the Leucegene samples using the published truth set (Figure 4B). The TCGA-LAML sensitivities are computed using the callers default-filters while both strategies are shown for the Leucegene samples. Since only 58 SNVs are available in the Leucegene truth set, the same SNVs detected across downsampled runs are used to compute the sensitivity curves (see ‘Computation of sensitivity’ section in Methods).

**Figure 4:**
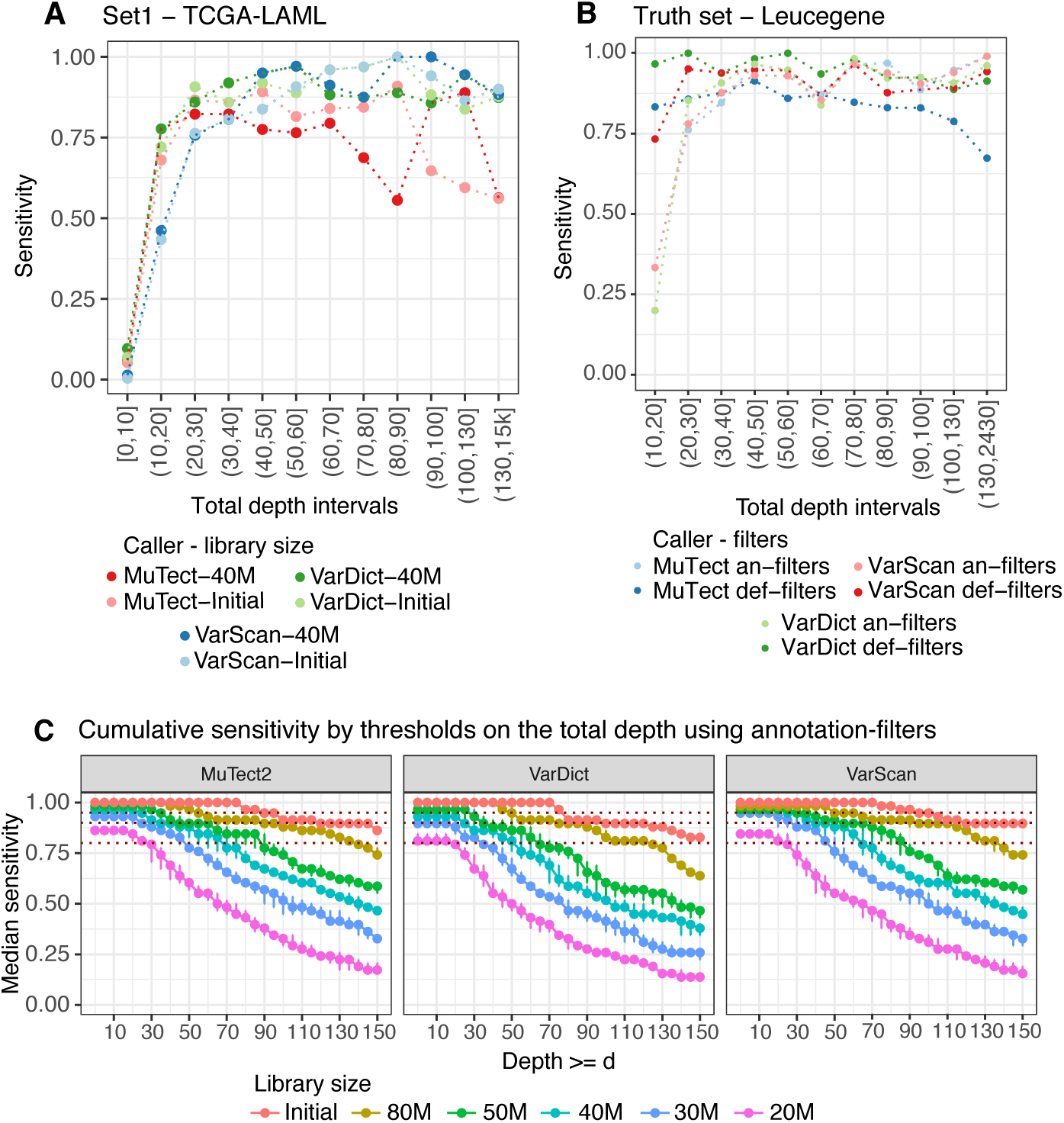
Sensitivity by total depth at a variant site. **A**. Sensitivity as a function of the total depth at a variant site using the initial and 40M TCGA-LAML libraries and adopting callers with default-filters. **B**. Sensitivity as a function of the total depth at a variant site using the Leucegene samples. The sensitivity is computed using the variants in the truth set, combining the calls from all downsampling runs, and using both types of filters. **C**. Median with maximum and minimum sensitivity in recovering the SNVs in the truth set using the Leucegene samples. Only SNVs with total depth ≥ d are considered as called. The sensitivity by depth is computed for each starting library size (colours) and using annotation-filters. Each estimated median sensitivity (and minimum and maximum) is the median across random downsampling runs at the same library size. The red dotted lines represent the 80%, 90% and 95% sensitivity thresholds.

Across all strategies and in both datasets, the changes in sensitivities stabilise and remain above 75% when the total depth is larger than 20X, and they stay on average above 80% when larger than 30X. The low sensitivities obtained with the TCGA-LAML samples using the variants in Set1 are due to 67% of the variants having total depth in RNA below 20X (60% between 0-10X, 7.4% between 10-20X, Figure 4A, Figure S10) as the truth set was obtained from DNA variants. The difference in sensitivity between the initial and the downsampled TCGA-LAML libraries is small at any total depth interval, with the exceptions of MuTect, delivering lower and more discordant sensitivities at higher depths. When stratifying the Leucegene sensitivities by library size, the majority of the variants lost derives from the 20M and 30M libraries, using either filtering strategies (Figure S11). This is because the Leucegene variants were detected from RNA and are therefore well expressed. Indeed, all variants in the truth set have total depth >90X in the initial libraries (Figure S12).

Figure 4C shows the cumulative sensitivity of each caller using the annotation-filters, and applying increasingly higher thresholds, *d*, on the total read depth at a variant site. Only variants with total depth ≥ *d* are classified as called. The annotation-filters were used to allow a fair comparison with MuTect due to the bias with its default filters. However, the two strategies have a comparable performance (Figure S13). The results confirm the poor performance when using only 20M fragments. Indeed, using this library size the sensitivity remains below 90% at any total depth threshold. Not until the 40M library sizes, and borderline with 30M, can MuTect and VarScan recover at least 95% of the variants when requiring *d* ≥ 20. VarDict recall is slightly lower than the other callers, recovering 90% of the variants when setting *d* ≥ 20. When increasing the threshold *d* above 20, the calls are progressively more discordant between different library sizes.

## Discussions and conclusions

Our study complements previous research which focused on the mean-on-target depth necessary to recover SNPs detected from RNA-Seq, WES and WGS where different depths at a variant site, ranging between 10X and 40X were identified depending on the technology used[8, 7]. Previous work has suggested 20M PE reads is enough for accurate detection of DE genes[6], but our results suggest higher levels of coverage are required to permit robust variant detection. Among previous research, there is a lack of studies focusing on the total library size required for accurate detection of mutations in cancer transcriptomes, where variants occur at different VAF and which stand as a cost-efficient choice, especially when studying AML genomes. This paper is the first study analyzing the reduced detection of SNVs from bulk cancer RNA-Seq with progressively smaller library sizes. Our study provides a direct connection between the on-site variant features (VAF and total depth) and the total library size needed when planning the sequencing design.

We suggest that between 30M and 40M fragments are required to guarantee 90-95% sensitivity in recovering variants in myeloid genes, which is approximately 50-100% greater than the suggested library size for DE analysis. While the largest loss in sensitivity is often observed when sequencing less than 30M fragments, it is not clear how to define a fixed library size suitable for all types of samples. This is because of the incremental decrease in sensitivity as the library size gets smaller. Nonetheless, using the Leucegene samples, we saw a negligible reduction in the sensitivity to recover validated variants when sequencing 80M compared to more than 100M PE reads (Figure 1). Following this, there are similar modest losses of sensitivity when going from 80M to 50M, from 50M to 40M, and from 40M to 30M reads. The loss only becomes more noticeable with library sizes below 30M fragments. A similar conclusion was reached with the TCGA-LAML cohort, where the sensitivity in recovering SNVs on recurrently mutated myeloid genes remains almost unchanged when sequencing 40M rather than 60M PE reads (Figure 3C). We also found a total depth at a variant site larger than 20X to be a critical threshold to stabilize the sensitivity above 75%, for both AML cohorts (Figure 4A). However, the highest recall rates are obtained for variants with total depth larger than 50X.

Taking all the above results together, we can consider a general formula to inform the target library size, based on the expression level (RPKM) of key genes and the total depth at a variant site:

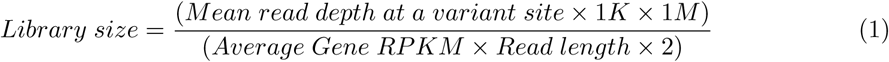

For example, if we want the average total depth at a variant site to be 30X, in order to ensure good sensitivity in a gene with RPKM of 4 (slightly above the lowest expression levels of genes in Set2, Figure 2B), and we consider 100 bp PE reads, the required library size is 37.5M fragments. If we set the total depth to 20X, then the library size drops to 25M fragments. By using the above equation and knowledge about the transcriptome features of the tissue under investigation one can explore whether detecting mutations from RNA-Seq is worthwhile.

We showed that the sensitivity in recovering validated SNVs in the Leucegene RNA-Seq samples is independent of the caller used, with the exception of MuTect with default settings, whose behavior should be adjusted when applied to RNA-Seq. However, the choice of a caller has a greater impact when calling indels, and targeted approaches are required to guarantee a good sensitivity. The number of bioinformatics tools developed recently to improve indel detection in RNA-seq is a demonstration of the increasing interest in exploiting RNA for variant calling, and the need for better algorithms and benchmarking[17, 18]. In particular, AML genomes often harbour hot spot indels whose detection is of clinical importance. These are internal tandem duplications (ITDs) found in FLT3 and KIT[19, 20] as well as a 4bp insertion in NPM1. The km algorithm was recently published[15] which performs targeted variant detection. The sensitivity and precision of the caller were studied using FLT3-ITDs and NPM1 insertions detected from the Leucegene and the TCGA-LAML cohorts, reaching more than 90% sensitivity for both lesions and making it an appealing caller for complex indels.

In this study, no single method to call and filter SNVs always outperformed the others. Prioritizing cancer variants based on external databases, like COSMIC, rescued NRAS somatic mutations which were found at very low VAF in one sample in the reference PON. NRAS is commonly mutated in CBF-AML and it is not surprising if the same mutation is present with low VAF in normal haematopoietic stem cells. The hard thresholds used in combination with the annotation-filters appear too stringent and induced sharper decreases in sensitivities compared to using default-settings in the Leucegene samples. Also, while MuTect and VarDict are better choices to detect complex indels (Figure 1A right plot), VarScan is quicker than MuTect; it allows genome-wide calls; and has comparable sensitivity to the other callers in recovering SNVs. A suggested workflow when calling variants in cancer RNA-seq would be to choose a SNV caller and adapt its filters to the specific characteristics of the cohort and samples available. For example, it is advisable to carefully check for highly recurrent filters to avoid losing interesting variants as was found with the clustered events filter in MuTect. In general, MuTect should not be used in tumor-only mode but this does raise the problem of the availability of suitable matched normal samples for tumour RNA-Seq. Differences in gene expression between tissues may complicated variant detection, and with blood cancers it can be challenging to obtain a normal sample, free from contaminating cancer cells. If normal DNA is available, it can be used to filter artefacts and germline variants[21, 22]. It is also useful to adopt different callers to detect different types of variants, e.g. using a targeted caller for indels to increase sensitivity and a fast genome-wide caller for SNVs.

In conclusion, this study offers a starting point to developing cost-efficient sequencing designs for other genomic studies. As cancers are extremely heterogeneous, a rigorous assessment of the cohort characteristics is necessary before making decisions for other tumour types. Also, this analysis was performed on diagnostic samples with a median of 60% bone marrow blast (= tumour cells) content, and it is advisable to consider adjusting the library size to account for tumour purity, in order to preserve detection power.

## Methods

### Availability of the data and materials

The FASTQ files of the 45 Leucegene CBF-AML RNA-Seq samples were downloaded from GEO at the accession numbers GSE49642, GSE52656, GSE62190, GSE66917, and GSE67039. The FASTQ files of the 17 RNA-Seq samples of CD34+CD45 RA-cord blood cells used to create the PON variants were downloaded from GEO at accession number GSE48846. The TCGA-LAML BAM files were downloaded after permission was granted from the GDC Data Portal at https://portal.gdc.cancer.gov/projects/TCGA-LAML[16]. The software settings and supporting scripts used for downsampling, preprocessing, alignment, variant calling and annotations are available on GitHub at https://github.com/annaquaglieri16/Supporting-scripts-library-size-RNA-Seq.

### Leucegene CBF-AML RNA-Seq: Alignment and pre-processing

The R package GEOquery[23] and the SRA Toolkit[24] were used to download the SRA files from GEO and to convert them to FASTQ files. FastQC[25] 0.11.5 was used to check the quality of the initial FASTQ files and no samples was removed due to low quality. Sample SRX381851 was excluded from the analysis due to a small initial library size of only 44.5M PE reads. The GNU Parallel command-line utility[26] was used to parallelize the FastQC runs. The FASTQ files were aligned against the UCSC hg19 reference genome to resemble the analysis performed in the original publication[9]. Alignment was performed with STAR[27] 2.5 in two-pass mode. The splice junctions from the 45 CBF-AML samples collected from the first pass were used to inform the alignment in the second pass. The reference fasta genome and gtf files necessary to build the STAR index were downloaded from https://sapac.support.illumina.com/sequencing/sequencing_software/igenome.html. STAR was chosen for several reasons: its speed; its good performance in the correct alignment of indels[28]; and since it is the suggested choice in the GATK[29] Best Practices for RNA-Seq variant calling. Read groups were added to the aligned BAM files using AddOrReplaceReadGroups from Picard tools[30] 2.9.4 and PCR duplicates were marked with sambamba[31] 0.6.6 markdup. Duplicate reads were not removed from the BAM files but reads marked as duplicates are ignored at the variant calling step. The quality of the BAM files were validated with ValidateSamFile from Picard Tools and no errors were found in any processed library. Gene counts were obtained with featureCounts using the hg19 inbuilt RefSeq annotation available in Rsubread[32]. The same pipeline was used for both the initial and every downsampled run as well the PON samples.

### TCGA-LAML data: bamfile pre-processing

The TCGA-LAML cohort comprises 151 50bp PE RNA-Seq bamfiles of which 17 are CBF-AML. The bamfiles were already aligned to the hg38 genome reference genome using STAR in two-pass mode. Read groups had already been added using STAR. We flagged PCR duplicates with sambamba markdup and obtained gene counts with featureCounts and the hg38 inbuilt RefSeq annotation. Both sambamba markdup and MarkDuplicates from Picard Tools failed in processing sample TCGA-AB-2931 which was removed from the rest of the analysis. Eight bamfiles failed the downsampling step due to some internal features of the bamfiles. These samples are: TCGA-AB-2821, TCGA-AB-2870, TCGA-AB-2884, TCGA-AB-2925, TCGA-AB-2950, TCGA-AB-2991, TCGA-AB-2994, and TCGA-AB-2995 and they were excluded from the rest of the analysis. Out of the 142 bamfiles left, 139 had mutations validated through targeted capture and manual review in the original publication[16]. After excluding indels from the truth set, 136 RNA-Seq samples with available variants were used for sensitivity analysis.

### Downsampling strategy

The FASTQ files from the Leucegene CBF-AML data were downsampled using the seqtk toolkit for FASTA/Q files[26]. Every fixed downsampled library size was obtained five times (only 3 times for the 80M library size). Five seeds were used to allow reproducibility of the results: 100, 26880, 56745, 7234, 9999. Only BAM files were available for the TCGA-LAML cohort and they were downsampled only once using the DownsampleSam function from Picard Tools. This function extracts a proportion of the reads out of the initial library size. We adjusted the sampling proportions for the TCGA-LAML samples in order to simulate a setting with approximately 40M sequenced fragments and a >90% mapping rate. Adjustment was needed since the mean proportion of mapped reads in the TCGA-LAML samples was lower then for the Leucegene samples, defining a ratio of 1.24 (93.4% in the Leucegene and 75.6% in TCGA samples respectively). Therefore, we first obtained the total number of reads in each TCGA-LAML BAM file using samtools flagstat[33], where this number includes both mapped and unmapped reads, and increased the sampling proportion of each sample by 1.24. This adjustment should make the results comparable and it is based on the assumption that, in the future, the mapping quality for bulk RNA-Seq samples will more likely resemble the Leucegene quality. This led to a mean of 37.3M mapped fragments in the downsampled TCGA-LAML cohort (37.4M mapped with the Leucegene data). Table S4 contains the downsampling proportions used for the TCGA-LAML RNA-Seq samples.

### TCGA truth sets

The table of validated variants was downloaded from https://tcga-data.nci.nih.gov/docs/publications/aml_2012/SupplementalTable06.tsv and was used as truth set for the sensitivity analysis with the TCGA-LAML data. From this table we removed 6,460 mutations falling outside of gene bodies; 8,319 variants in untranslated and intronic regions; and only kept variants belonging to samples whose RNA-Seq BAM files were available. Eventually, 1,643 SNVs were kept for sensitivity analysis (Table S5). From these SNVs, two truth sets are defined: Set1 includes all 1,643 SNVs; Set2 is a subset of Set1 including the published significantly mutated myeloid genes[16], out of which FAM5C and HNRNPK are removed by previous filters, and on top of which 7 genes are added as they are mutated in the Leucegene CBF-AML samples[9] (KMT2C, JAK2, GATA2, CSF3R, ASXL1, NF1, KDM6A) (Set2 genes reported in Table S3). We created 1k symmetric windows around the starting positions of the variants in the final truth set and we used those regions to perform variant calling. The genomic position obtained from the original table were lifted up from the hg18 to the hg38 reference genome using the UCSC Genome Browser[34]. When matching the TCGA-LAML variants called by a caller with the variants in the truth sets, we did not use transcript information. This was because the transcripts reported in the original table derive from old annotations as well as they come from a mix of ensembl and RefSeq annotations. We decided to not consider transcript information to avoid wrong assignments or removal of variants due to missing transcript information.

### Choice of callers and BAM file preparation for variant calling

Several variant callers are compatible with RNA-Seq[35]: RADIA[21], Seurat[22], SNPiR[36], eSNV-detect[37], VarScan[12] and VarDict[13]. The first two callers, RADIA and Seurat, integrate tumor-normal RNA and DNA and were not considered since only RNA is available. SNPir is a caller specifically developed for RNA-Seq but for normal tissue. It is based on GATK[29] pre-processing and the Haplotypecaller[38] and implements a series of RNA-Seq specific filters. eSNV-detect implements an ensemble approach by combining the calls performed with SAMtools using two different aligners. Here we compared the performance of three popular callers: VarScan 2.4.0 (which requires samtools mpileup[33] output), VarDict 1.5.1 and MuTect2 from GATK 3.7.0, which is the GATK choice for tumor samples. Only VarDict was developed to call variants from both RNA and DNA. VarDict can call variants in both tumor-only and matched tumor-normal settings, whereas VarScan and MuTect were designed for somatic variant calling. All three callers were run in tumor-only mode and a reference of PON samples was created for filtering (see details ‘Variant filtering’ section). Prior to calling variants with MuTect and VarScan the BAM files were pre-processed following the GATK best practices, which include splitting reads that contain N’s in the CIGAR string and base quality recalibration. VarDict can handle spliced reads without pre-processing. VarDict can only call variants on subsets of the genome whereas both VarScan and MuTect call variants genome-wide. This last difference will not introduce any bias into our comparison since variants are called only in target regions of interest throughout the whole analysis. The three callers call both SNVs and indels but only VarDict and MuTect adopt local realignment, which should give better accuracy around indels. Samtools mpileup performs base alignment quality (BAQ) which aims at reducing SNVs miscalled due to nearby indels.

### Variant calling, annotation and standardised output

To allow a fair comparison between variant callers, variants were called with MuTect, Samtools mpileup + VarScan and VarDict using their default settings. Variant calling is always performed in tumour-only mode. Variants were then annotated with the Variant Effect Predictor (VEP) 89.0[39]. The genome assemblies GRCh37 and GRCh38 were used for the Leucegene and the TCGA-LAML samples respectively. The annotated VCF files were parsed using the parsing functions included in the varikondo[40] R package to produce a standardized output across callers containing the relevant information for the analysis.

### Variant filtering

We adopted two variant filtering strategies, namely default-filters and annotation-filters. The default-filters strategy is based on each caller’s default filters while the annotation-filters strategy exploits external databases and applies the same quality measures across callers. The current analysis aims at reproducing a general framework to be used in tumor-only RNA-seq variant calling, and we do not wish to “overfit” a caller’s settings to benefit this specific type of data. What matters in this study is to compare the performance of variant calls under the same circumstances while varying the overall library size.

### Default filters

We used only variants flagged with PASS in the VCF output to estimate sensitivity and specificity based on each caller’s default settings. We also used a PON samples to remove likely artefact variants that are called across many normal independent samples. In particular, we used a set of 17 RNA-Seq libraries of CD34+CD45RA-cord blood cells from 17 non-pooled individuals. We created a PON variants separately for every caller (see details in ‘Variant filtering’ section in Additional Methods). A variant is classified as present in normals if it is found in more than two normal samples or in less than two normal samples but with VAF > 0.03. We created PON variants using both the hg19 and hg38 reference human genomes.

### Annotation filters

We set up a series of filters based on variant quality measures, public databases, features of the genome known to be challenging (repeat regions, homopolymers, splice junctions), and PON samples. Some of these filters, including variants in homopolymers, RNA editing sites[41], variants in repeat regions[42] and variants near splice junctions, have been previously shown to be successful in reducing the number of false positives[36, 41]. Variant annotation using VEP adds information from external databases of mutations such as COSMIC[10], ExAC[43] and dbSNP[44]. Details about how to download the databases mentioned here and how we used them to flag variants can be found in section ‘Variant filtering’ in Additional Methods. Flags and quality filters are then used for filtering. In particular:

- Variants are removed if they are found in the PON (using the same strategy as explained in the ‘default-filters’ section) and they are not found in COSMIC.
- Variants are removed if they are not found in COSMIC but they are present in dbSNP and ExAC.
- Even if present in COSMIC, variants are removed if they overlap with exon boundaries, homopolymer stretches or repeated regions. Overlap with exon boundaries is defined if the variant falls within 4bp upstream of an exon start site or downstream of an exon end site. Exon boundaries were obtained from the hg19 and hg38 inbuilt RefSeq Rsubread annotation. We considered as homopolymers, stretches of the same nucleotide longer than 5bp as obtained from the hg19 and hg38 reference human genomes (more details provided in ‘Annotation databases and genomic features’ section in Additional Methods).
- Finally, a variant is kept only if it has an average base quality > 18, a minimum alternative allele depth of 5, a minimum total depth of 15 and a minimum VAF of 0.03. This means, for example, that in order to keep a rare variants with VAF < 0.03 the mutation needs to be covered by at least 167 reads.

### Matching alternative alleles

In both strategies a variant is classified as called in a downsampled library only if the alternative allele matches the alternative allele in the truth set, if available, or the one obtained when calling variants using the initial deeply sequenced libraries. The alternative alleles were available for the TCGA-LAML dataset but not for the Leucegene cohort. The three callers were consistent in reporting the same alternative alleles for all the SNVs called with the Leucegene cohort but not for indels. To reduce callers differences in reporting indels, we first ran the variant normalization tool vt normalize[45]. To take into account persisting differences we considered a match if an indel lied within 100bp (+/-50bp) of the position reported in the Leucegene truth set. Manual curation was then needed to remove false positives. An extra level of complexity derives from the fact that callers report composite events differently, making it hard to assess a match with the alternative allele. MuTect and VarScan output indels as separate SNVs and short indels rather than as block substitutions, while VarDict, like the km algorithm, outputs composite variants in the same line. This is why we also explored whether a partial match with the reference and alternative alleles would increase sensitivity for MuTect and VarScan. A partial match required at least 3bp overlap with the alleles reported in the truth set.

### Computation of sensitivity

At every downsampled run, the *sensitivity* was computed as in Equation 2 below:

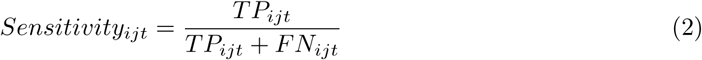

Where *TP*_*ijt*_ is the number of variants called by caller *i* with library size *j* that belong to truth set *t*, and *FN*_*ijt*_ is the number of variants in truth set *t* missed by caller *i* at the library size *j*. A variant is reported as a match with respect to a truth set, if it is called by a caller and it matches the genomic information (chromosome, position etc..) reported in the truth set.

The *sensitivity by intervals on total depth* is computed as in Equation 2 but stratifying by intervals on the total depth. The intervals are set from 0X to the maximum total depth observed in the data by gaps of 10 until 100X. When total depth >100X only two intervals are considered, 100X-130X and 130X to the maximum total depth.

The sensitivity as a function of the depth *d* is computed as in Equation 3 below:

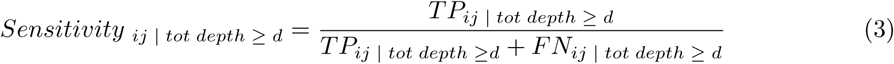

Where *TP*_*ij* | *tot depth ≥ d*_ is the number of variants called by caller *i* with library size *j* that has total depth *≥ d* and that belong to the truth set. *FN*_*ij* | *tot depth ≥ d*_ is the number of variants with total depth *≥ d* present in the truth set but missed by caller *i* with library size *j*.

### R packages used in the study

The variant calling workflow was developed using the package optparse[46]. The packages foreach[47] and doParallel[48] were used to parallelize the parsing of the variant annotation fields added by VEP. Variants output were standardised across callers using the package varikondo[40], available on GitHub. The Bioconductor package GenomicRanges[49] was used to create the files needed to annotate variants with respect to genomic features (see ‘Additional Methods’ in Additional File 1 for more details). The library seqinr[50] was used to read FASTA files into R in order to detect stretches of homopolymers used for variant filtering. The package samplepower[51], only available on GitHub, contains the functions used to compute sensitivities throughout the analysis (more details in ‘Additional Methods’, Additional File 1). Data manipulation to parse variants output and to produce summary of the sensitivity results was obtained using the R packages readr[52], dplyr[53], tidyr[54], stringr[55]. All figures in this paper were produced with the libraries ggplot2[56] and cowplot[57]. All analysis in R were run using R3.5.2.

## Additional files

### Additional file 1

Supplementary document containing supplementary figures, tables and methods. (PDF 4Mb)

## Author’s contributions

AQ performed all the analyses and wrote the manuscript. CF provided guidance through several steps of the development of the RNA-Seq variant calling workflow and reviewed the manuscript. IM conceived the initial idea and provided suggestions and revision for the manuscript. TS contributed with suggestions during the conception and development of the study and reviewed the manuscript.

## Acknowledgements

This study was supported by the Melbourne International Research Scholarship (AQ); in part by the NHMRC Program Grant 1054618 (TPS); by grants from the Australian National Health and Medical Research Council (Project Grant to IJM 1145912; Independent Research Institutes Infrastructure Support Scheme grant 9000220), the Cancer Council Victoria (grant-in-aid to IJM 1124178), a Victorian State Government Operational Infrastructure Support (OIS) grant; a Victorian Cancer Agency fellowship (to IJM) and the Felton Bequest.

The results here are based upon data generated by the Leucegene consortium (https://leucegene.ca/) and the TCGA Research Network: http://cancergenome.nih.gov/.

