## Additional File 1 for "Finding a suitable library size to call variants in RNA-seq"

### Additional files

Anna Quaglieri

### 1 Additional figures

#### 1.1 Figure S1. Frequency of mutated AML genes in the Leucegene cohort

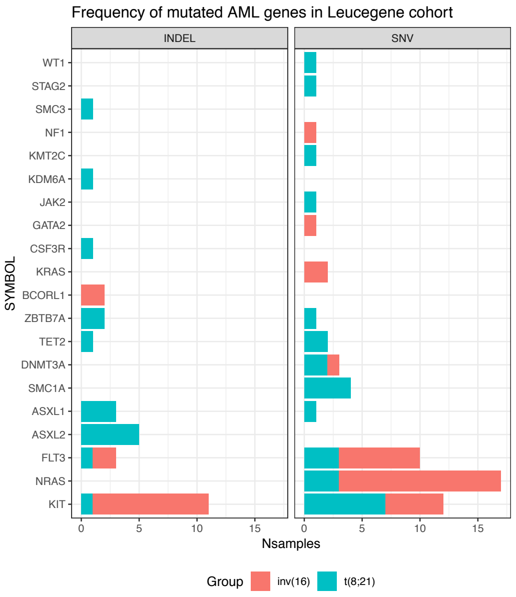

**Figure S1:** Number of SNVs and indels in the original publication using the Leucegene CBF-AML RNA-Seq samples[1]. Frequency is provided by gene and by CBF subtype.

#### 1.2 Figure S2. Flags used in MuTect with default filters

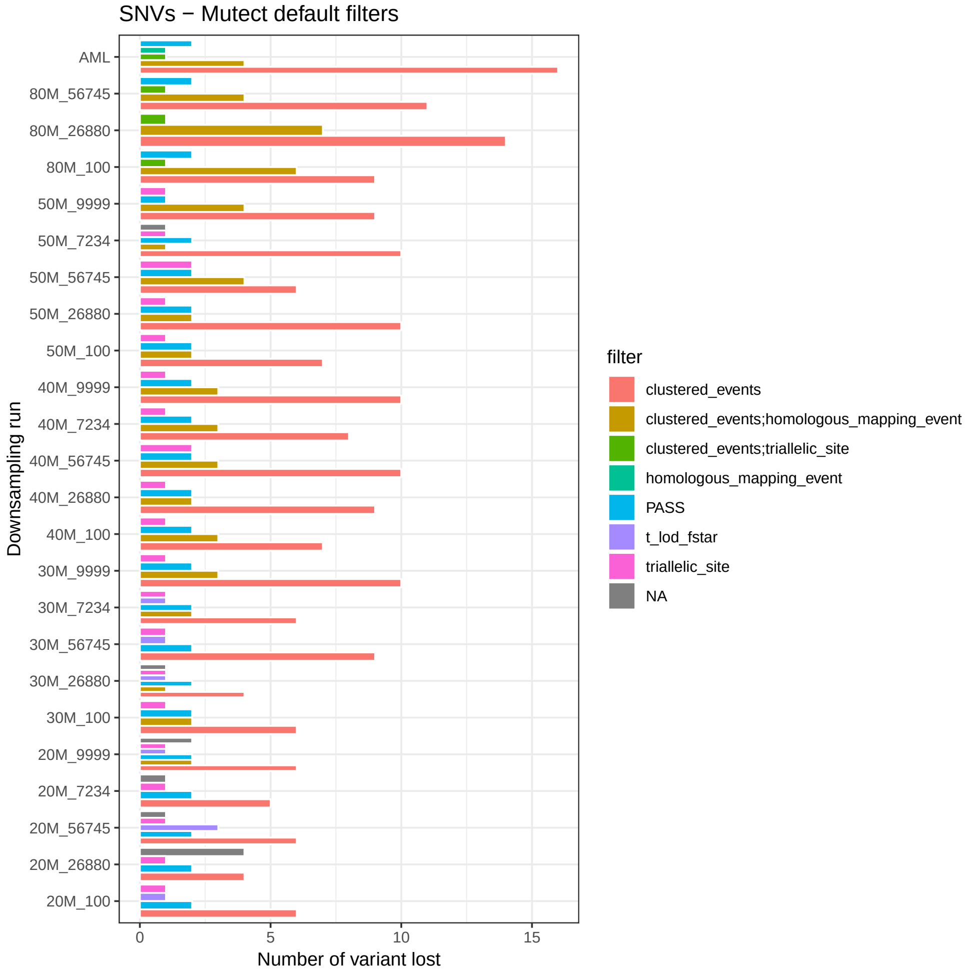

**Figure S2:** The barplot shows the breakdown of the filters used by MuTect to filter variants at every downsampling stage. The filter is `NA` if a variant from the truth set was not present in the MuTect's VCF output. The `clustered_events` filter is responsible for the loss of the majority of the variants and for the strange behaviour of MuTect, which improves its sensitivity at shallower depths. This is an issue already reported for this caller (`https://github.com/broadinstitute/gatk-protected/issues/773`) and also previously observed[**Coudray2018-yw**] where the `clustered_events` filtered the largest numbers of variants in RNA-Seq samples. The thickness of the bars is due to the different number of flags reported across downsampled runs but it has no meaningful interpretation.

#### 1.3 Figure S3. Drop in sensitivity between subsequently smaller library sizes

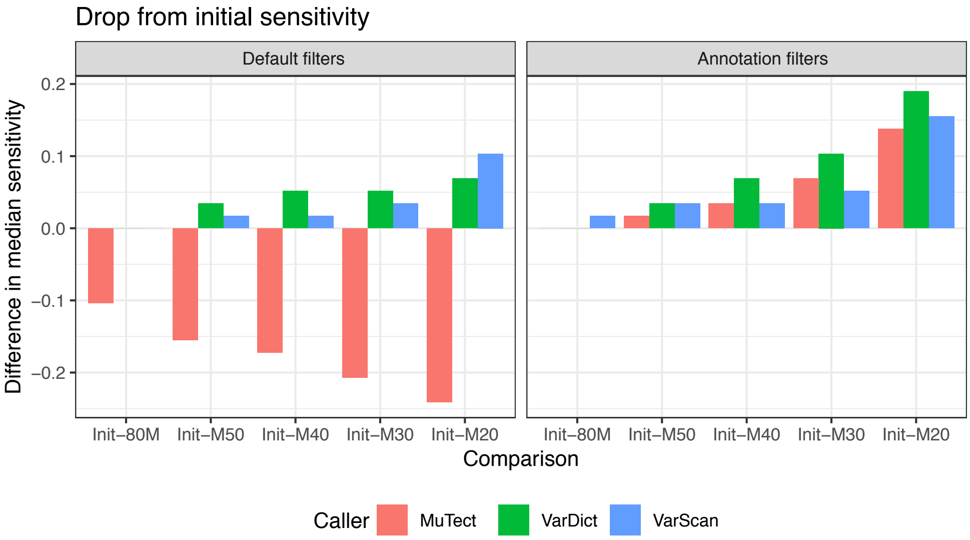

**Figure S3:** Differences in median sensitivity in recovering SNVs between the initial RNA-Seq libraries and each downsampled run. Differences are shown by caller and by filtering strategy.

#### 1.4 Figure S4. Indel sensitivity with the published truth set using partial match

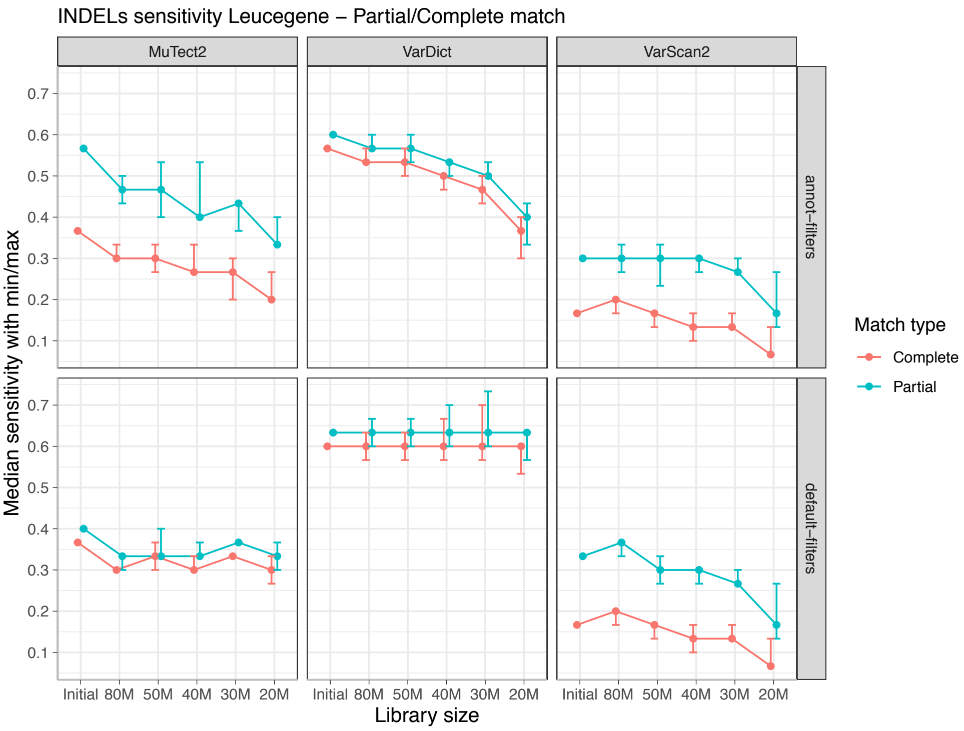

**Figure S4:** Median, maximum and minimum sensitivity (y-axis) of recovering the indels in the Leucegene truth set allowing for a partial (blue lines) or complete (red lines) match between the alternative allele reported by the caller and the one reported in the truth set.

#### 1.5 Figure S5. Flags of the variants missed by each caller using the published truth set

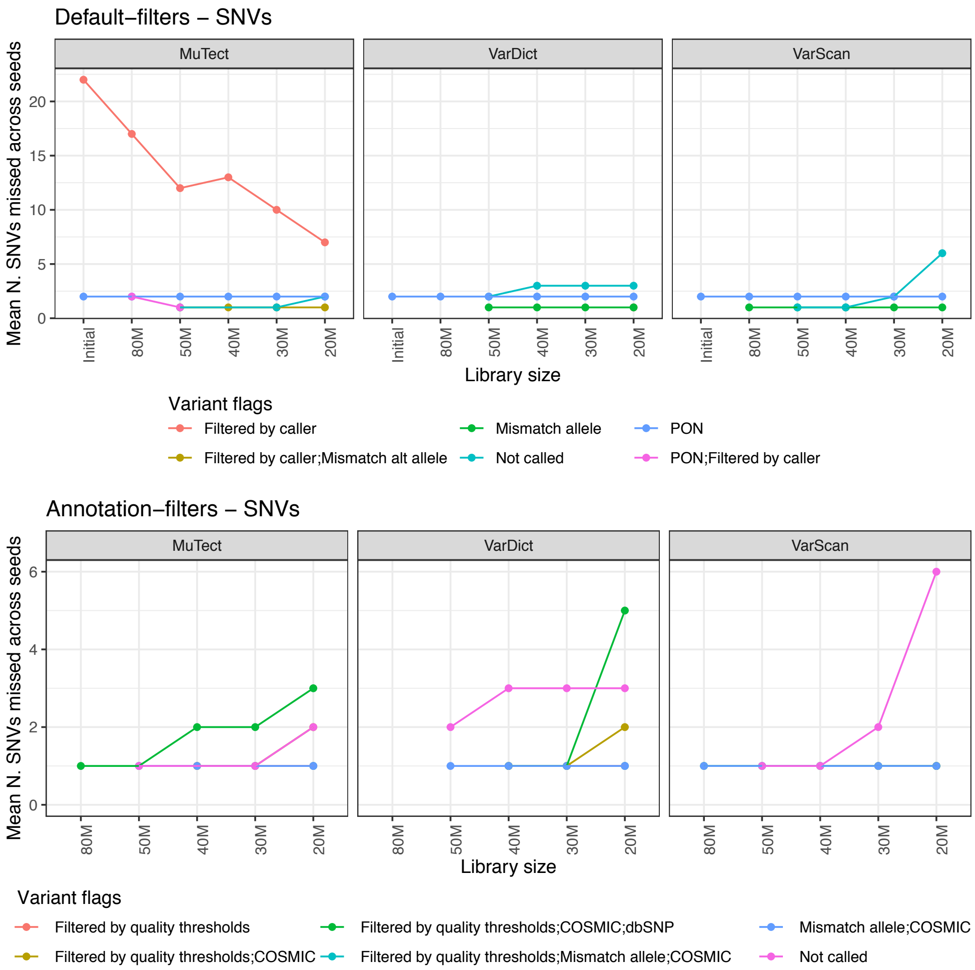

**Figure S5:** Lineplots showing the flags of the SNVs lost at decreasing the library size, using default-filters (top panel) and annotation-filters (bottom panel). The flags are: *Filtered by caller* if the FILTER field in the VCF output by a caller was different from PASS; *PON* if present in the normals; *Mismatch allele* if there is a mismatch between the alternative allele in the downsampled set and the truth set; *Not called* if it wasn't in the VCF produced by a caller; *Filtered by quality thresholds* if it didn't pass the quality thresholds reported in section 'Annotation filters: filter variants using external databases' in Methods; *dbSNP* if present in the dbSNP database; *COSMIC* if present in the COSMIC database of cancer mutations. There is one COSMIC variant (blu line bottom panel) which is filtered out by the three callers also at the 50M and 80M library sizes. This is variant chr4:55599340 from sample SRX729623. This is a cKIT activating mutation (N822K) annotated as a T $->$ G mutation in `https://cancer.sanger.ac.uk/cosmic/mutation/overview?id=1322`. This variant is called in six samples in this cohort. In four samples the mutation called is a T $->$ A substitution and in two samples a T $->$ G substitution. Sample SRX729623 harbours both variant alleles but with different VAFs and the three callers only report the alternative allele with the highest VAF. In some of the random downsampling runs these frequencies are inverted and the other alternative allele is detected.

#### 1.6 Figure S6. Change in total depth at a variant site using smaller library sizes

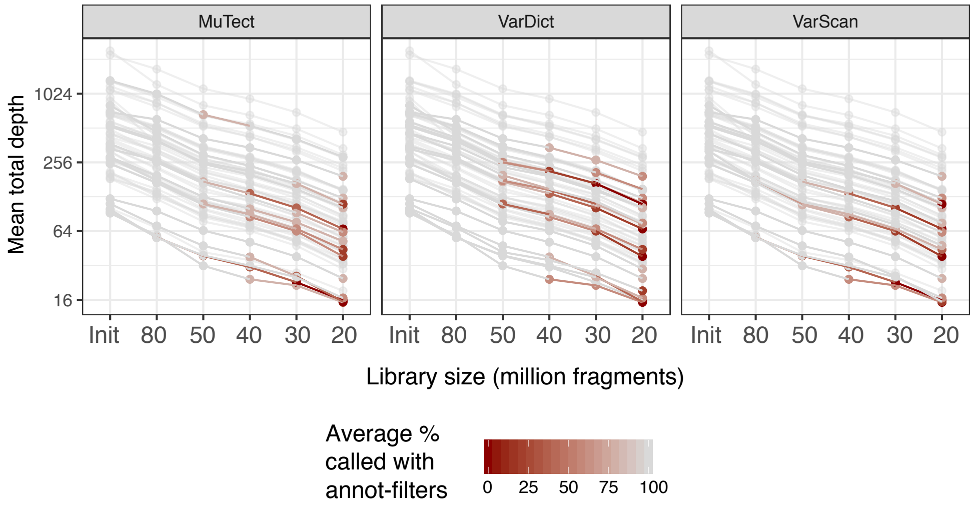

**Figure S6:** Total depth at a variant site on a logarithmic scale (y-axis) for the variants in the truth set using different library sizes (x-axis). Ticks on the y-axis are on the original scale. A line in the plots represents one mutation in the published truth set. Each dot is coloured according to the average number of times a variant was called by one caller using the annotation-filters across replicated downsampling runs at one specific library size. This should reflect the variability in random downsampling and the progressive loss of mutations as the library size gets smaller.

#### 1.7 Figure S7. Total versus alternative depth at a variant site for the variant in the Leucegene truth set

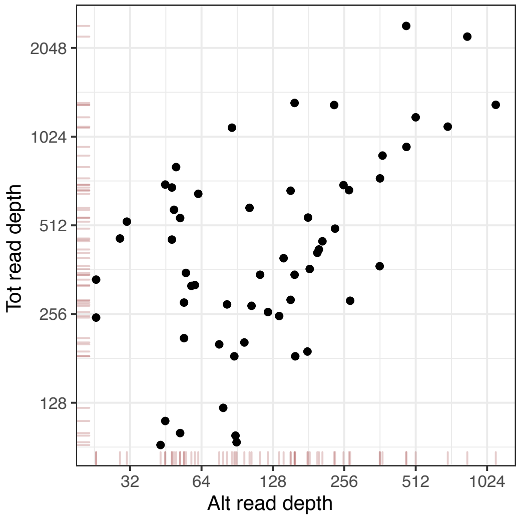

**Figure S7:** Total versus alternative depth at a variant site for the variant in the Leucegene truth set reported in Table 1. The x and y axis are on the logarithm scale.

#### 1.8 Figure S8. Flags of the variants missed using the TCGA-LAML cohort

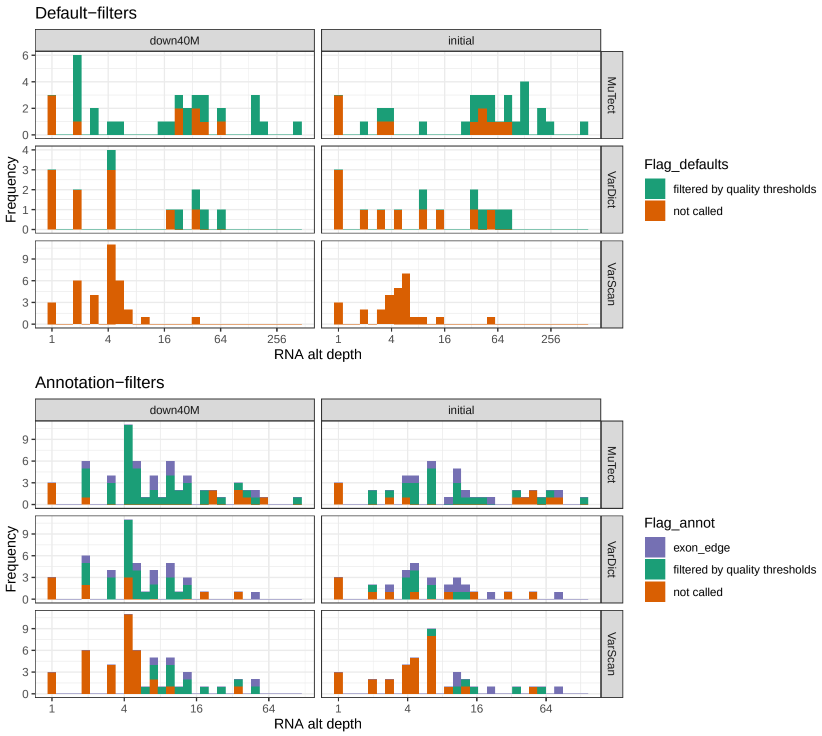

**Figure S8:** Flags of variants missed with the TCGA-LAML cohort using the initial and downsampled library sizes and using the default-filters (top plot) and the annotation-filters (bottom plot). In both cases the truth set includes all the variants in *Set2*, i.e. variants on recurrently mutated myeloid genes (Table S3).

#### 1.9 Figure S9. Sensitivity in recovering mutations on recurrently mutated AML genes using the TCGA-LAML cohort and annotation-filters

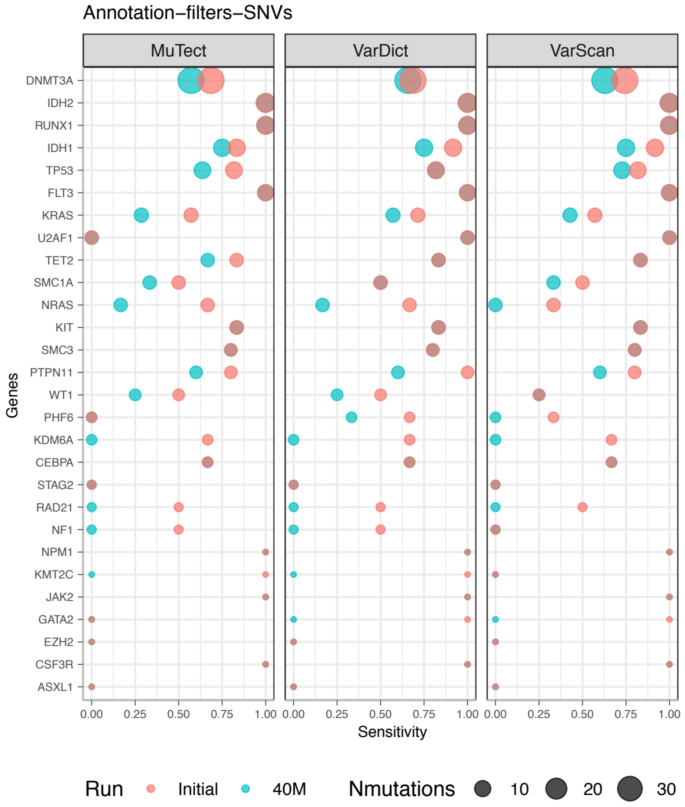

**Figure S9:** Sensitivity in recovering mutations on recurrently mutated AML genes (Set2) using the TCGA-LAML cohort and annotation-filters. The size of the dots is proportional to the number of times a gene is mutated and genes were ordered by mutation load, with the most mutated genes at the bottom. Red dots corresponds to the results obtained with the initial library sizes and blue dots using the 40M fragments libraries.

#### 1.10 Figure S10. Number of variants in each total depth interval using the TCGA-LAML samples

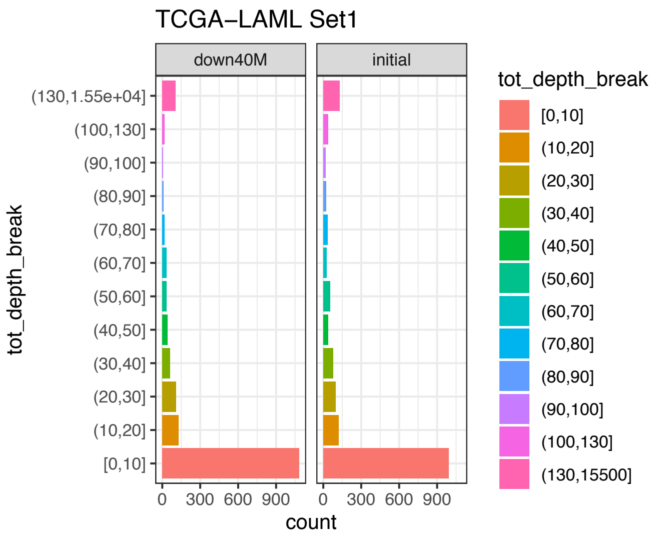

**Figure S10:** Number of variants present in each total depth interval using the TCGA-LAML variants from Set1 and stratifying by library size.

#### 1.11 Figure S11. Sensitivity by total depth in the Leucegene cohort stratified by caller and library size

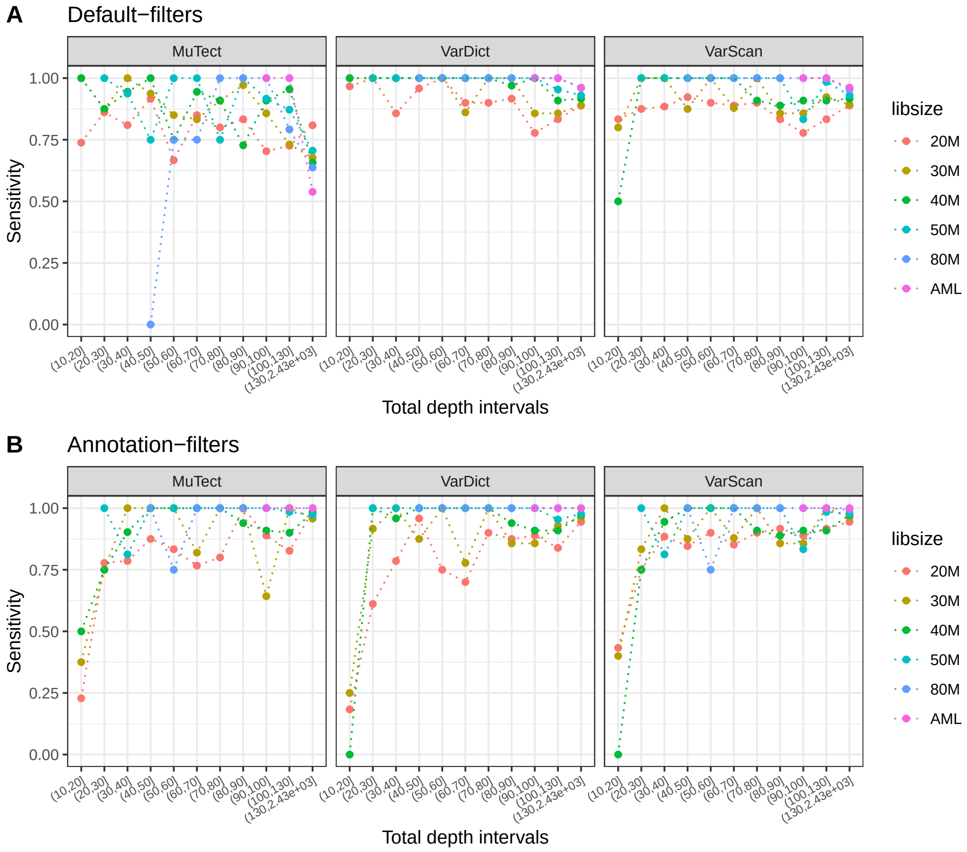

**Figure S11:** **A**. Sensitivity by interval on the total depth at a variant site using default-filters with the Leucegene samples. The sensitivity is stratified by callers (panels) and by total library size (colours). **B**. Same as **A** but using annotation-filters.

#### 1.12 Figure S12. Number of variants in each total depth interval using the Leucegene samples

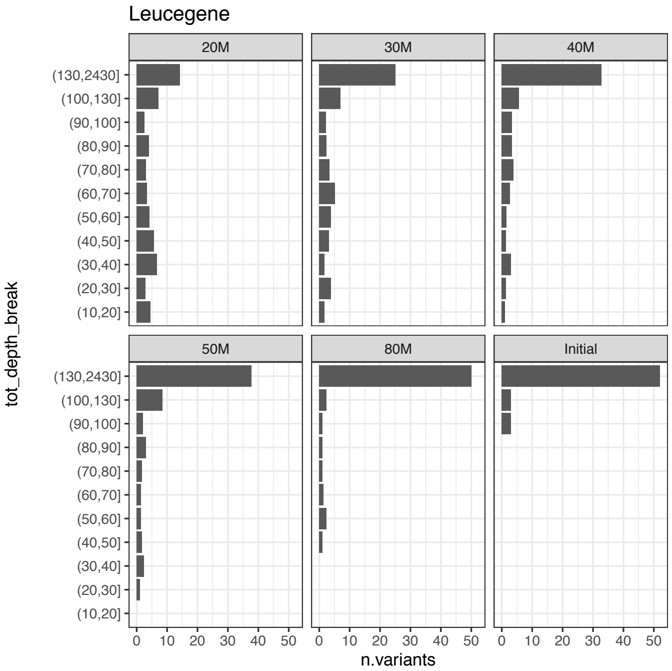

**Figure S12:** Number of variants present in each total depth interval using the Leucegene samples and their validated truth set of variants, stratifying by library size.

#### 1.13 Figure S13. Sensitivity by total depth at a variant site and by total library size

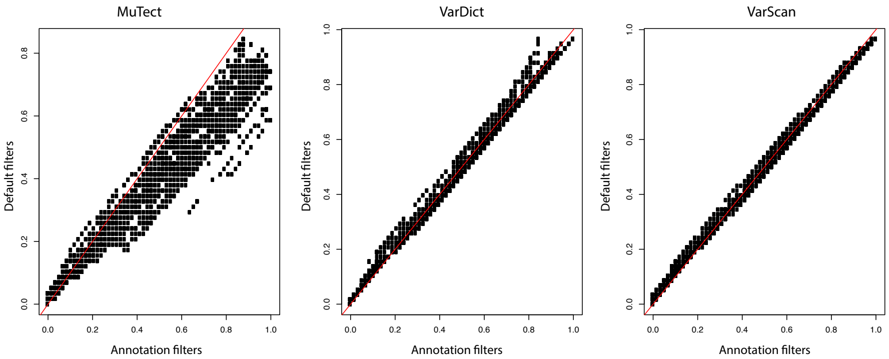

**Figure S13:** Mean sensitivity in recovering the SNVs in the *truth set*, considering variants with total depth $\geq$ d across different starting library sizes. The scatterplot compares the sensitivity obtained with the default-filters against the annotation-filters across all the thresholds used. The annotation-filters are consistently more sensitive than the default-filters for MuTect. The red line represents the identity line.

### 2 Additional tables

#### 2.1 Table S1. Gene regions used to call variants for the sensitivity analysis with the Leucegene sample

| **Chr** | **Start** | **End** | **Gene** | **Strand** |
|---|---|---|---|---|
| chr1 | 36932144 | 36948415 | CSF3R | - |
| chr1 | 115247585 | 115259015 | NRAS | - |
| chr10 | 112326949 | 112364892 | SMC3 | + |
| chr11 | 32409822 | 32456581 | WT1 | - |
| chr12 | 25358223 | 25403365 | KRAS | - |
| chr13 | 28577911 | 28674229 | FLT3 | - |
| chr17 | 29421445 | 29705195 | NF1 | + |
| chr19 | 4045716 | 4066316 | ZBTB7A | - |
| chr2 | 25456330 | 25564959 | DNMT3A | - |
| chr2 | 25962753 | 26100812 | ASXL2 | - |
| chr20 | 30945647 | 31027622 | ASXL1 | + |
| chr3 | 128198765 | 128211530 | GATA2 | - |
| chr4 | 55523595 | 55607381 | KIT | + |
| chr4 | 106066532 | 106201460 | TET2 | + |
| chr7 | 151832510 | 152132590 | KMT2C | - |
| chr9 | 4984745 | 5128683 | JAK2 | + |
| chrX | 44731921 | 44972357 | KDM6A | + |
| chrX | 53401570 | 53449177 | SMC1A | - |
| chrX | 123093910 | 123237005 | STAG2 | + |
| chrX | 129138664 | 129192558 | BCORL1 | + |

**Table S1:** Genomic regions used to call variants for the sensitivity analysis with the Leucegene sample.

#### 2.2 Table S2. Characteristics of the mutations lost by each caller at different library sizes.

| Size | Caller | N | medVAF | minVAF | maxVAF | medAltD | medTotD | medBlast |
|---|---|---|---|---|---|---|---|---|
| 20M | MuTect | 8 | 0.2 | 0 | 1 | 5 | 38 | 65 |
| 20M | VarDict | 10 | 0.1 | 0 | 1 | 6 | 39 | 63 |
| 20M | VarScan | 9 | 0.1 | 0 | 1 | 5 | 39 | 65 |
| 30M | MuTect | 4 | 0.2 | 0.02 | 1 | 7 | 61 | 73 |
| 30M | VarDict | 5 | 0.1 | 0.02 | 0.89 | 7 | 98 | 58 |
| 30M | VarScan | 3 | 0.1 | 0.02 | 1 | 6 | 57 | 58 |
| 40M | MuTect | 2 | 0.2 | 0.03 | 1 | 7 | 71 | 61 |
| 40M | VarDict | 4 | 0.1 | 0.03 | 0.45 | 9.5 | 140 | 56 |
| 40M | VarScan | 2 | 0.25 | 0.03 | 1 | 7 | 78 | 61 |
| 50M | MuTect | 1.5 | 0.5 | 0.03 | 1 | 31 | 92 | 71 |
| 50M | VarDict | 2.5 | 0.1 | 0.03 | 0.26 | 17 | 182 | 71 |
| 50M | VarScan | 2 | 0.5 | 0.03 | 1 | 22.5 | 71 | 69 |
| 80M | MuTect | 1 | 1 | 1 | 0.96 | 52 | 54 | 81 |
| 80M | VarScan | 1 | 0.6 | 0.1 | 1 | 40 | 115 | 73 |

**Table S2:** Summary of the number and characteristics of the mutations lost at different library sizes. From left to right the table reports the total library size fragments (Size); the caller used (Caller); the mean number of variants missed across replicates downsampled to the same library size (N); the median, minimum and maximum VAF of the variants missed (medVAF, minVAF, maxVAF); the median and total depth of the alternative allele (medAltD, medTotD); and the median bone marrow blast content percentage of the samples whose mutations were lost (medBlast). These measures were obtained for every location directly from the bamfiles using the GATK `DepthOfCoverage` function. For example, a minimum VAF of zero means that among all the variants lost at a library size with a caller, there was at least one location for which no reads supporting the alternative allele were found.

#### 2.3 Table S3. Genes used to define *Set*2 in the TCGA-LAML sensitivity analysis

The list in Table S3 is an intersection of the significantly mutated myeloid genes defined in the original TCGA-LAML paper[2] and the set of genes mutated in the Leucegene CBF-AML patients[1].

| Symbol | Symbol |
|---|---|
| FLT3 | SMC3 |
| NPM1 | PHF6 |
| DNMT3A | STAG2 |
| IDH2 | RAD21 |
| IDH1 | FAM5C |
| TET2 | EZH2 |
| RUNX1 | HNRNPK |
| TP53 | ASXL1 |
| NRAS | ASXL2 |
| CEBPA | BCORL1 |
| WT1 | CSF3R |
| PTPN11 | GATA2 |
| KIT | JAK2 |
| U2AF1 | KDM6A |
| KRAS | KMT2C |
| SMC1A | NF1 |
| ZBTB7A | |

**Table S3:** List of genes used to define Set2 in the sensitivity analysis using the TCGA-LAML RNA-Seq samples.

#### 2.4 Table S4. Downsampling proportions for the TCGA-LAML samples

**Table S4:** Sample-specific proportions used to downsample the TCGA-LAML samples.

#### 2.5 Table S5. TCGA-LAML truth set

**Table S5:** SNVs in Set1 used as truth set for the sensitivity analysis using the TCGA-LAML cohort. Set2 can be obtained by subsetting the table considering only genes in Table S3.

### 3 Additional methods

#### 3.1 Extract depth at a variant site from the bamfiles

The GATK function `DepthOfCoverage` was used to recover the VAF, the total depth, the depth of the alternative allele of the locations in the truth sets for every downsampled run directly from the bamfile.

```
module load gatk/3.7.0
gatk -T DepthOfCoverage \
-R genome.fa \
-o ./sample_loci_depth_of_coverage \
-I sample.bam \
--printBaseCounts \
-L ./mutations_loci.bed
```

#### 3.2 Variant filtering

Below is the code used to create the annotation information used to filter variants. The code was run with R3.5.2. The R package `samplepower`[3], available on GitHub, was developed to combine all the functions used to analyse the variant calling outputs in this paper and to compute sensitivity and specificity measures across callers.

#### 3.3 Panel of normals (PON)

The 17 CD34+ RNA-Seq samples were processed using the same pipeline used for the cancer samples. The chromosome and position reported by a caller were used to filter variants found in the tumour samples. Therefore, no annotation with VEP was needed. A PON was created for every caller, combining the variant calls of the 17 CD34+ RNA-Seq samples using the GATK functions `CombineVariants` and `VariantsToTable`. PON variants were created after aligning the 17 RNA-Seq samples to the hg19 and the hg38 reference genomes, to allow filtering for the Leucegene CBF-AML (hg19) and the TCGA-LAML (hg38) data respectively.
`VariantsToTable` extracts fields of interest from the a `VCF` file. The settings used were the same for all callers expect from VarScan which required: `set -GF FREQ` to be set instead of `set -GF AF` due to a different way of specifying the variant allele frequency in the VCF. Below is the code used.

```
module load gatk/3.7.0
module load vcflib/1.0.0-rc1
gatk -T CombineVariants \
-R path_hg19_reference/genome.fa \
--variant:SRX322282 normal_cd34_caller/SRX322282_germline_snvs_indels.vcf \
--variant:SRX322283 normal_cd34_caller/SRX322283_germline_snvs_indels.vcf \
```

```
--variant:SRX322284 normal_cd34_caller/SRX322284_germline_snvs_indels.vcf \
--variant:SRX322285 normal_cd34_caller/SRX322285_germline_snvs_indels.vcf \
--variant:SRX322286 normal_cd34_caller/SRX322286_germline_snvs_indels.vcf \
--variant:SRX322287 normal_cd34_caller/SRX322287_germline_snvs_indels.vcf \
--variant:SRX322288 normal_cd34_caller/SRX322288_germline_snvs_indels.vcf \
--variant:SRX322289 normal_cd34_caller/SRX322289_germline_snvs_indels.vcf \
--variant:SRX322290 normal_cd34_caller/SRX322290_germline_snvs_indels.vcf \
--variant:SRX322291 normal_cd34_caller/SRX322291_germline_snvs_indels.vcf \
--variant:SRX322292 normal_cd34_caller/SRX322292_germline_snvs_indels.vcf \
--variant:SRX322293 normal_cd34_caller/SRX322293_germline_snvs_indels.vcf \
--variant:SRX322294 normal_cd34_caller/SRX322294_germline_snvs_indels.vcf \
--variant:SRX322295 normal_cd34_caller/SRX322295_germline_snvs_indels.vcf \
--variant:SRX322296 normal_cd34_caller/SRX322296_germline_snvs_indels.vcf \
--variant:SRX322297 normal_cd34_caller/SRX322297_germline_snvs_indels.vcf \
--variant:SRX322298 normal_cd34_caller/SRX322298_germline_snvs_indels.vcf \
-o normal_cd34_caller/PON_caller_target_regions.vcf \
-L ./Leucegene_target_genes.bed \ # restrict to regions of interest
-genotypeMergeOptions UNIQUIFY
gatk  -T VariantsToTable \
-R path_hg19_reference/genome.fa \
-V normal_cd34_caller/PON_caller_target_regions.vcf \
--splitMultiAllelic \
-F CHROM -F POS -F ID -F REF -F ALT -F FILTER -F set -GF AF \
-o normal_cd34_caller/PON_caller_target_regions.table
```

The `samplepower::parsePON()` R function is then used to: read the table of normal variants into R; summarising in how many samples a variant was called and what is the minimum VAF. The VAF field is parsed depending on the caller used. This information are needed for filtering variants found in cancer samples. The `caller` arguments is one of `varscan`, `mutect` or `vardict`.

```
# Example with VarScan
var_linkPON <- file.path("PON_varscan_target_regions.table")
var_pon <- samplepower::parse_pon(var_linkPON, caller = "varscan")
var_pon$Location <- paste(var_pon$CHROM, var_pon$POS, sep = "_")
```

#### 3.4 Annotation databases and genomic features

**COSMIC, dbSNP and ExAC** We used VEP annotations to define if a variant was present in the COSMIC[4], dbSNP[5] and ExAC[6] databases.

**RNA editing sites** We downloaded the database of RNA editing sites for the hg19 human reference genome from `http://rnaedit.com/download/` and processed it in R to convert it to a `GRanges`[7] object. We used the UCSC Genome Browser to liftover the RNA editing sites from hg19 to hg38 to be used with the TCGA-LAML samples.

```
library(GenomicRanges)
RADAR_hg19_allRNAedit <- read_table2("RADAR_hg19_allRNAedit.txt",
col_names = FALSE)
colnames(RADAR_hg19_allRNAedit) <- c("chrom", "start", "end")
RADAR_hg19_allRNAedit <- subset(RADAR_hg19_allRNAedit, chrom %in% chroms)
RADAR_hg19_allRNAedit <- subset(RADAR_hg19_allRNAedit,
!is.na(RADAR_hg19_allRNAedit$start) &
    !is.na(RADAR_hg19_allRNAedit$end))
RNAeditGR <- GRanges(RADAR_hg19_allRNAedit)
```

**Repeated regions** We downloaded the RepeatMasker track from the UCSC Genome Browser for the hg19 and hg38 human reference genome, saved it into a `.bed` file and converted it into `GRanges` objects to be used with Leucegene and TCGA-LAML cohorts respectively.

```
library(GenomicRanges)
# Repetitive regions
hg19_repeat_regions <- read_table2("hg19_repeat_regions.bed",
col_names = FALSE)
colnames(hg19_repeat_regions) <- c("chrom", "start", "end",
"name", "X5", "strand")
hg19_repeat_regions <- subset(hg19_repeat_regions, chrom %in% chroms)
repeatsGR <- GRanges(hg19_repeat_regions)
```

**Exon boundaries** We removed variants falling within 4bp of an exon boundary to reduce false positives due to RNA splicing. We used the `hg19` (and `hg38` for the TCGA-LAML cohort) `Rsubread` in-built Refseq gene annotation. We created genomic ranges containing 4bp upstream the start of an exon and 4bp downstream the exon end. The NCBI gene annotation, downloaded from `ftp://ftp.ncbi.nlm.nih.gov/gene/DATA/GENE_INFO/`, was used to match gene Symbols to entrez GeneIDs.

```
library(GenomicRanges)
library(Rsubread)
ncbi <- read.delim(gzfile(file.path("Homo_sapiens.gene_info_29-11-2017.gz")))
hg19 <- Rsubread::getInBuiltAnnotation(annotation = "hg19")
chroms <- c(paste0("chr", 1:22), "chrX", "chrY", "chrM")
match_symbol <- match(hg19$GeneID, ncbi$GeneID)
hg19$Symbol <- ncbi$Symbol[match_symbol]
hg19_exons_start <- GRanges(seqnames = hg19$Chr,
IRanges(start = hg19$
Start -
    4,
end = hg19$Start), strand = hg19$
Strand,
mcols = data.frame(side = rep("left",
    nrow(hg19))))
hg19_exons_end <- GRanges(seqnames = hg19$Chr,
IRanges(start = hg19$End,
end = hg19$End +
    4), strand = hg19$
Strand, mcols = data.frame(side = rep("right", nrow(hg19))))
hg19_exons_combined <- append(hg19_exons_start, hg19_exons_end)
```

**Homopolymers** The code below was used to create the `GRanges` object of homopolymers (stretched of 5bp with the same nucleotide). The hg19 and hg38 UCSC reference genome FASTA files were downloaded from `http://sapac.support.illumina.com/sequencing/sequencing_software/igenome.html`.

```
library(seqinr)
library(GenomicRanges)
chroms <- c(paste0("chr", 1:22), "chrX", "chrY", "chrM")
fastaHg19 <- read.fasta("ref_genome.fa")
fastaHg19_chr <- fastaHg19[names(fastaHg19) %in% chroms]
# Make an RLe for every chromosome
rle_Hg19 <- lapply(fastaHg19_chr, function(chrom) {
    Rle(as.numeric(as.factor(chrom)))
```

```
})
# for every element of an RLE for every chromosome save the start, the end,
# and the length of the rle run
rle_Hg19_homop <- lapply(rle_Hg19, function(chrom) {
    end = end(chrom)
    start = start(chrom)
    len <- runLength(chrom)
    data.frame(start, end, len) %>% filter(len > 5)
})
# Combine them all to create a GRanges object
rle_chrom <- do.call(rbind, rle_Hg19_homop)
rle_chrom$chrom <- rep(names(rle_Hg19_homop), times = sapply(rle_Hg19_homop,
    nrow))
GRanges_homop <- GRanges(seqnames = rle_chrom$chrom,
IRanges(start = rle_chrom$start,
    end = rle_chrom$end))
```

##### 3.5 Annotate variants and compute sensitivity with samplepower

The R function `samplepower::variant_power()` is the main function of the package and it computes sensitivity with respect to a truth set. Within this function: 1) the variants called from the downsampled and initial RNA-Seq are loaded 2) the variants are then flagged with information about the annotation databases and the PON variants discussed above, using the function `samplepower::flag_variants()`. Based on the annotation and variants quality measures, discussed in the Methods section 'Variant filtering', each variant present in the truth set is classified as called/not called at any downsampling run. After the variants are flagged, the coverage and VAF information provided with the output from GATK3 `DepthOfCoverage` are merged to the variants. The merging is achieved by the `samplepower::parse_gatk_coverage()` function. At this stage, only SNVs present in the truth sets (both for the Leucegene and the TCGA-LAML cohorts) are analysed, and the sensitivity is computed. The recovery of indels starts with finding indels called by a caller within a region around the genomic locations of indels in the truth set (Methods section 'Matching alternative allele'). However, the final decision about whether an indel is called will require manual curation and the final sensitivity is computed after curation.
